# Interneuron diversity in the human dorsal striatum

**DOI:** 10.1101/2023.03.22.533839

**Authors:** Leonardo D. Garma, Lisbeth Harder, Juan M. Barba-Reyes, Mónica Díez-Salguero, Alberto Serrano-Pozo, Bradley T. Hyman, Ana B. Muñoz-Manchado

## Abstract

Deciphering the striatal interneuron diversity is key to understanding the basal ganglia circuit and to untangle the complex neurological and psychiatric diseases affecting this brain structure. We performed single-nucleus RNA-sequencing (snRNA-seq) of postmortem human caudate nucleus (CN) and putamen (Pu) samples to elucidate the diversity and abundance of interneuron populations and their transcriptional structure in the human dorsal striatum. We propose a new taxonomy of striatal interneurons with eight main classes. We provide specific markers for all subclasses and validated some of them with quantitative *in situ* fluorescence hybridization, such as a novel PTHLH-expressing population that exhibits different abundance and gene expression between CN and Pu. For the most abundant interneuron populations in human striatum, PTHLH and TAC3, we found matching known mouse interneuron populations based on key functional genes such as ion channels and synaptic receptors. Remarkably, human TAC3 and mouse Th populations share important similarities including the expression of the neuropeptide tachykinin 3. Finally, we were able to integrate our dataset with several prior smaller human striatal snRNA-seq studies, thus supporting the generalizability of this new harmonized taxonomy.

## Introduction

The dorsal striatum is a subcortical brain structure that in human consists of caudate nucleus (CN) and putamen (Pu), separated by the internal capsule. Together with the ventral striatum (nucleus accumbens and olfactory tubercle), the globus pallidus, the subthalamic nucleus, and the substantia nigra, it makes up the basal ganglia nuclei^1^. The striatum carries out functions related to motor control, action learning, reward-related behavior, and cognition with certain regional preferences. The CN is mainly responsible for eye movement and cognitive functions, the Pu for motor control, learning and auditory responses, and the ventral striatum is related to limbic functions such as reward and motivation. Dysfunction of the striatum is a key feature of neurodegenerative disorders such as Parkinson’s and Huntington’s diseases^2,3,4,5^ as well as of psychiatric conditions such as obsessive-compulsive disorder and schizophrenia^6,7,8^.

The dorsal striatum is the main input area of the basal ganglia and exhibits a high level of activity-dependent synaptic plasticity^9^, representing a critical hub for the process and selection of information sent to the other basal ganglia nuclei. This information is relayed through the projecting neurons, known as medium spiny neurons (MSNs) because of their morphological features^10^. MSNs, which are characterized by their inhibitory signaling via gamma-aminobutyric acid (GABA), constitute the majority of the striatal neuronal population. However, their function depends on a diverse group of locally-projecting neurons known as interneurons.

Striatal interneurons integrate incoming information from different brain areas and act on MSNs activity to modulate the output information. This filtering process is also regulated by incoming dopaminergic and serotonergic projections from the midbrain and the dorsal raphe nucleus, respectively^11,12^. The aspiny striatal interneurons have been classically differentiated into two main groups: a small group of cholinergic giant neurons and a diverse population of GABAergic medium size neurons, based on a variety of specific markers and electrophysiological profiles^13,14,15^. Because the striatal interneurons have received little attention compared to the MSNs, consensus regarding the populations comprising these neuronal groups and how to identify them is lacking. However, recent advances such as new transgenic reporter mice that target to the complete striatal and cortical interneuron repertoire^15,16^, and single cell/nucleus RNA-sequencing (sc/nRNA-seq) have enabled large-scale approaches to investigate cell diversity based on the individual cell transcriptome^17,18,19^ in different mouse brain areas including the striatum^20,21,22^. Using these methods, a recent study identified seven interneuron populations in the mouse striatum based on their molecular and electrophysiological profile: Npy/Sst, Npy/Mia, Cck/Vip, Cck, Chat, Th, and Pthlh^20^. The *Pthlh*-expressing interneurons represent a novel class on striatal interneurons that is characterized by a variable *Pvalb* expression level and a broad continuum of intrinsic electrophysiological properties, which correlates with *Pvalb* levels^20^. This continuum seems to follow a regional gradient pattern within the mouse dorsal striatum suggesting that the different types of striatal cells receive inputs from different brain cortical areas^23^.

However, interneuron diversity in the human CN and Pu in terms of abundance and molecular identity remains unsolved. Most of the studies are limited by the technical approach because they have relied on the classical markers to identify interneuron populations and focused primarily on the cholinergic cells, expressing choline acetyltransferase (ChAT)^24,25,26^. Prior snRNA-seq studies on the human and non-human primate striatum have highlighted different aspects, such as broad differences across species and brain areas^27,28^ or in health vs. disease^29^, but lack sufficient interneuron sampling to characterize striatal interneuron diversity. In the present study, we have used snRNA-seq to investigate the diversity of interneurons on the human dorsal striatum (CN and Pu) from a total of 28 donors comprising nearly half a million nuclei overall, by far the largest study of this kind to date. We have leveraged this large dataset to establish the major and minor divisions between the interneuron classes and types in both regions and provide specific markers for each. We have validated part of our classification in tissue sections, confirming novel populations and markers (such as PTHLH, PVALB, and DACH1), identified differences between CN and Pu, and demonstrated an internal gradient structure within cell subclasses. Moreover, we have discovered key synapse-related genes in our taxonomy and described the consistency link between mouse and human PTHLH and TAC3 classes. Our taxonomy resisted the test of a comparison with prior human striatal snRNA-seq datasets, supporting efforts toward a new consensus classification of striatal interneurons.

## Results

### Interneuron heterogeneity in the human dorsal striatum

With the objective to further decode the diversity of interneurons in the human dorsal striatum, we isolated and sequenced single nuclei from fresh frozen CN (N = 25) and Pu (N = 28) samples of 28 neurotypical donors (Supplementary table 1). Samples were processed using an established snRNA-seq workflow, which allowed the enrichment of neuronal population by applying fluorescent-activated nuclei sorting. A subset of six Pu samples was additionally utilized in high-sensitivity fluorescent *in situ* hybridization (FISH) experiments to validate RNA expression patterns in tissue sections (Figure 1A).

**Figure 1.**
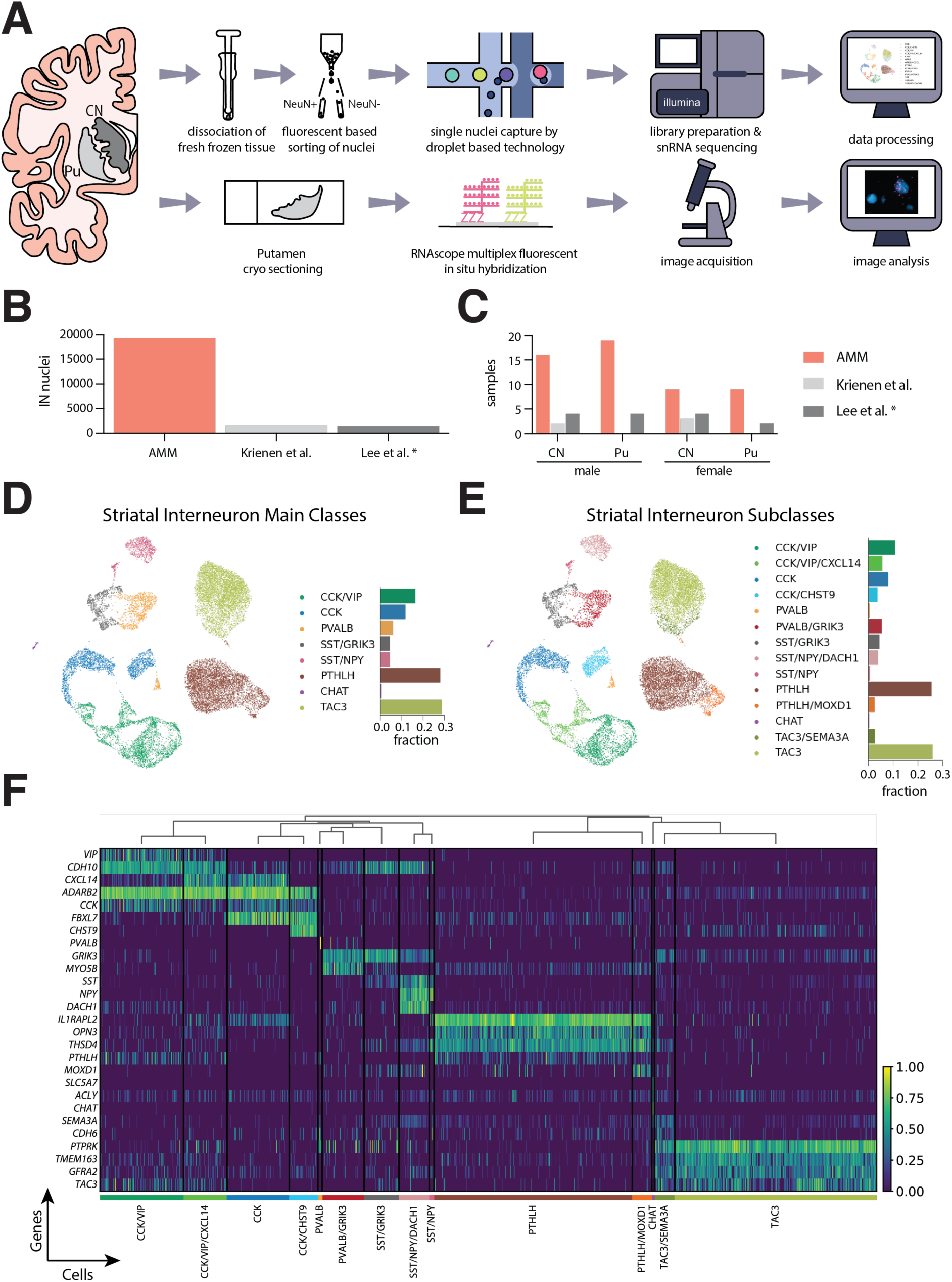
Interneuron heterogeneity of the human striatum determined by single-nucleus RNA sequencing. **A.** Schematic overview of the experimental design. **B.** Number of interneuron nuclei sequenced in the present study (labeled as AMM) vs. two other previous works. **C.** Number of human samples included in this study (AMM) vs. two previous works. **D**. UMAP projection of the snRNA-seq data of the nuclei labeled as interneurons, colored by interneuron class. The barplot next to the UMAP indicates the fraction of all interneuron nuclei represented by each class. **E**. UMAP projection of the interneuron nuclei colored by subclass. The barplot indicates the fraction of each subclass over all interneuron nuclei. **F**. Heatmap showing the expression of selected marker genes for each of the fourteen interneuron subclasses identified. The expression of each gene is normalized by its maximum value across all nuclei. The dendrogram above the heatmap indicates the proximity across subclasses based on the average Pearson correlation coefficient across all genes between each pair of subclasses. * Note that here only samples from control donors from Lee’s study are shown. CN, caudate nucleus; IN, interneuron; Pu, putamen.

The sequencing yielded 455,886 nuclei, out of which we discarded 29.4% after a quality control pipeline (Supplementary figure 1). From the remaining nuclei, we selected the interneurons through an iterative classification process in which we discarded glial cells, MSNs, and excitatory neurons based on bona-fide markers—astrocytes (*AQP4*, *ADGRV1*), microglia (*CSF1R*, *FYB1*), oligodendrocytes (*MBP*, *MOG*, *MAG*), oligodendrocyte precursor cells (*PTPRZ1*, *PDGFRA*, *VCAN*), vascular cells (*EBF1*, *ABCB1*, *ABCA9*), MSNs (*PPP1R1B*, *DRD1*, *DRD2*), and excitatory neurons (*SLC17A7*)—and selected positively for nuclei expressing *GAD1* and/or *GAD2*. This classification process resulted in **19,339** nuclei labeled as interneurons, representing the largest dataset of the human dorsal striatal interneurons available to date. The interneuron populations represented **10.67 %** of the total neuronal cells.

After clustering the data, we identified eight main interneuron classes: CCK/VIP (*ADARB2*+, *CCK*+, and *VIP*+), CCK (*ADARB2*+ and *CCK*+), PVALB (*PVALB*+), SST/GRIK3 (*SST*+ and *GRIK3*+), SST/NPY (*SST*+ and *NPY*+), PTHLH (*PTHLH*+ and *OPN3*+), CHAT (*CHAT*+ and *SLC5A7*+) and TAC3 (*TAC3*+ and *PTPRK*+) (Figure 1B), which could be divided into fourteen different subclasses identified by unique transcriptomic patterns (Figure 1C, D). The complete results of a differential expression analysis at class and subclass levels are provided in Supplementary table 2.

In our dataset, PTHLH and TAC3 constitute the largest interneuron classes, accounting for 28% and 28.6% of all detected interneurons, respectively. Both PTHLH and TAC3 contained small subclasses, distinguishable from the main type by the expression of *MOXD1* and *SEMA3A*, respectively. The *ADARB2*+ population (CCK and CCK/VIP classes) was equally abundant (28.1% of all interneurons), but exhibited a higher heterogeneity, with four subclasses clearly differentiated by specific marker genes. Although we followed the classical division between CCK and CCK/VIP, we observed that the *ADARB2*+ neurons could also be divided by the expression of the chemokine ligand *CXCL14* and the cadherin *CDH10* (Supplementary figure 2). We also found a great diversity of transcriptomic profiles among the neurons expressing *PVALB* and *SST*, as these two classes could be divided into five different subclasses based on the expression levels of two novel marker genes: *GRIK3,* which encodes the Glutamate Ionotropic Receptor Subunit 3, and *DACH1,* which encodes the Dachshund Family Transcription Factor 1. These five subclasses together represent 15% from the total of interneurons.

Interestingly, we also found a smattering of TAC3 expression in the CCK and CCK/VIP populations, therefore the TAC3 population is best defined by its high expression level of *PTPRK* (protein tyrosine phosphatase receptor type K). Similarly, we also noted low levels of *OPN3* (Opsin3)—one of the marker genes of the PTHLH cells—in the CCK and CCK/VIP populations.

### Validation of interneuron taxonomy with fluorescent *in situ* hybridization

The magnitude of the snRNA-seq dataset produced in the present study allowed us to detect novel interneuron populations with distinct transcriptomic profiles (Figure 1C, D). Therefore, we sought to validate some of these subclasses through quantitative multiplex FISH using up to three marker genes. Using probes against *SST*, *NPY* and *DACH1*, we were able to detect cells double-positive for *SST* and *NPY* in the Pu of all six donors assayed; the same applied to cells triple-positive for *SST*, *NPY*, and *DACH1* (Figure 2B-D). This is in line with our sequencing data (Figure 2A) and supports our decision to split the main class of SST/NPY interneurons into the SST/NPY and SST/NPY/DACH1 subclasses. In addition, in this FISH we identified a group of cells that were only positive for *SST* and most likely correspond to the subclass SST/GRIK3. The most conspicuous group of cells in this FISH was only positive for *DACH1*, which cannot be explained by the low *DACH1* expression found in CCK/VIP and PVALB/GRIK3 subclasses alone (Figure 2A); however, an extended search for *DACH1* expression in our dataset revealed that MSNs as well as some astrocytes, endothelial cells, and pericytes are positive for this marker (data not shown).

**Figure 2.**
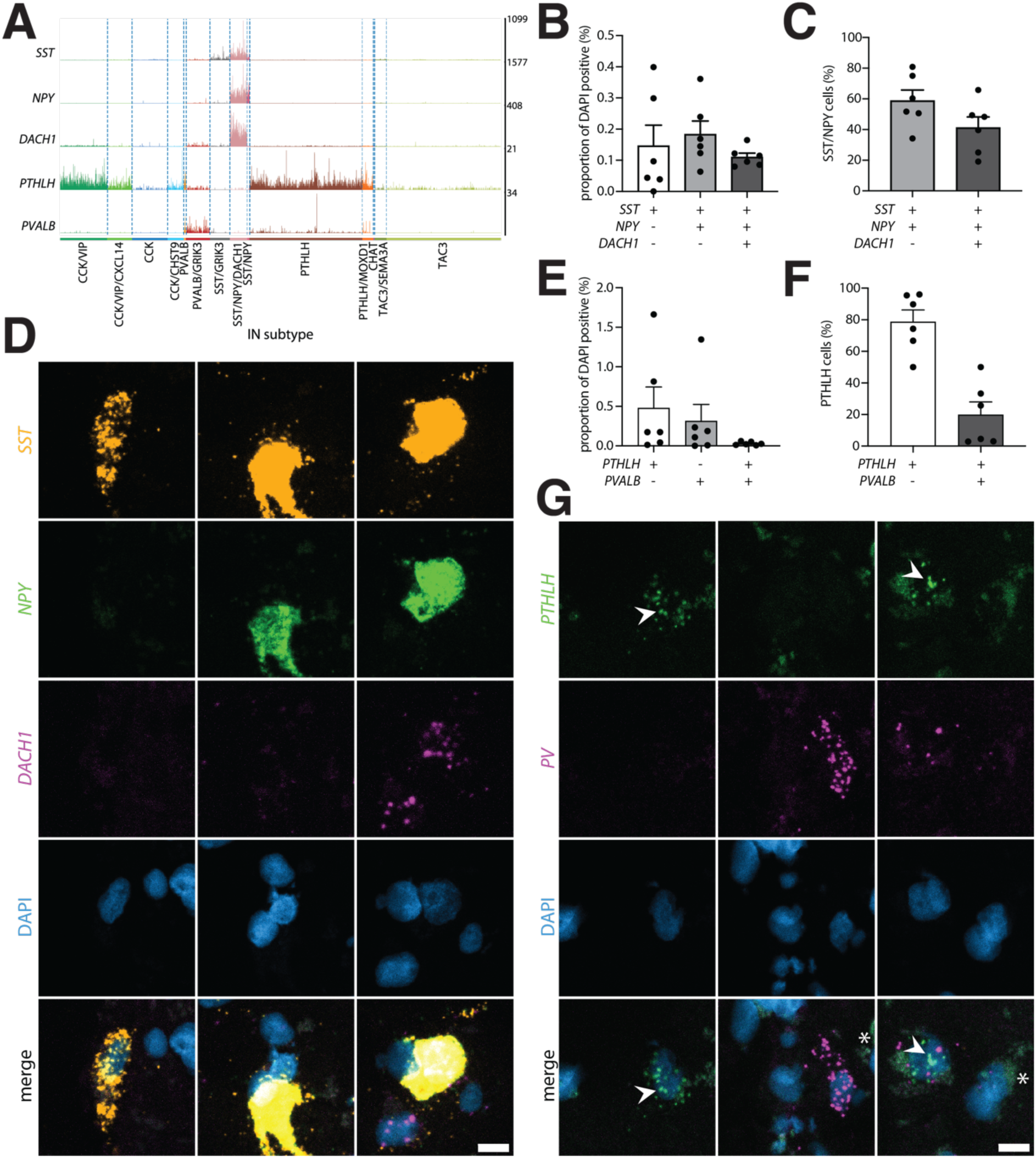
Validation using quantitative fluorescence *in-situ* hybridization confirms the existence of novel interneuron subclasses in the putamen. **A.** Tracks plot depicting raw UMI counts per nucleus for *SST, NPY*, *DACH1, PTHLH*, and *PVALB* in caudate and putamen. **B.** Proportion of cells positive for *SST*, *SST* and *NPY* or *SST*, *NPY*, and *DACH1* based on the total number of cells identified per donor (N = 6, putamen). **C.** Proportion of *SST* and *NPY* double-positive cells negative or positive for *DACH1* (N = 6, putamen). **D.** Representative images for (left) single-, (middle) double- and (right) triple-positive cells from the same donor, scale bar 10 µm. **E.** Proportion of positive cells for *PTHLH*, *PVALB*, or *PTHLH* and *PVALB* based on the total number of cells identified per donor (N = 6, putamen). **F.** Proportion of *PTHLH* positive cells negative or positive for *PVALB* (N = 6, putamen). **G.** Representative images for (left and middle) single- or (right) double-positive cells from the same donor. White arrows indicate spot signal for *PTHLH*, asterisks mark autofluorescence due to lipofuscin, scale bar 10 µm.

*PVALB*-expressing interneuron subclasses are of particular interest because PVALB has traditionally been used to identify a class of striatal interneurons. In contrast to the mouse striatum, in which *Pvalb*+ interneurons have been described to be contained within the *Pthlh*+ cells^20^ (i.e., all *Pvalb*+ interneurons are *Pthlh*+), our snRNA-seq data from human CN and Pu indicates the presence of a distinct *PVALB* positive but *PTHLH* negative subclass of interneurons. FISH using *PTHLH* and *PVALB* probes revealed *PTHLH* single-positive cells as well as *PTHLH* and *PVALB* double-positive cells in all six donors analyzed while *PVALB* single-positive cells were found in five of the six donors (Figure 2E). For both single-positive groups, the number of detected cells was highly variable across donors, while double-positive cells occurred in low numbers in all of them. Thus, while this experiment proves that *PTHLH* and *PVALB* double-positive cells exist in both human and mouse dorsal striatum, we provide evidence of a group of *PVALB* interneurons expressing minimal or no *PTHLH* at all in the human Pu and CN (Figure 2A and 2G).

### Interneuron populations exhibit region-based differences within the striatum

While we found all the interneuron classes identified in our snRNA-seq data in both the CN and the Pu (Figure 3A), we did note slight differences in abundance between both regions: CN was significantly richer in PTHLH interneurons (35.6% vs. 20.3%, p-value=0.001), whereas the CCK, SST/GRIK3 and PVALB classes were significantly more abundant in the Pu (12.1% vs. 7.1%, 3.5% vs. 2.4%, and 5.5% vs. 2.7%, respectively, with the belonging p-values in the same order 0.008, 0.027 and 0.015; Figure 3C). Several interneuron classes also exhibited distinct region-dependent transcriptomic signatures. Most notably, we found that the PTHLH class had significant differences in the expression of 276 genes by region (i.e.,126 upregulated and 150 downregulated in CN vs. Pu). SST/NPY, CHAT, TAC3 and CCK/VIP classes also showed expression differences in CN vs. Pu, but of smaller magnitude (Figure 3B, Supplementary table 3).

**Figure 3.**
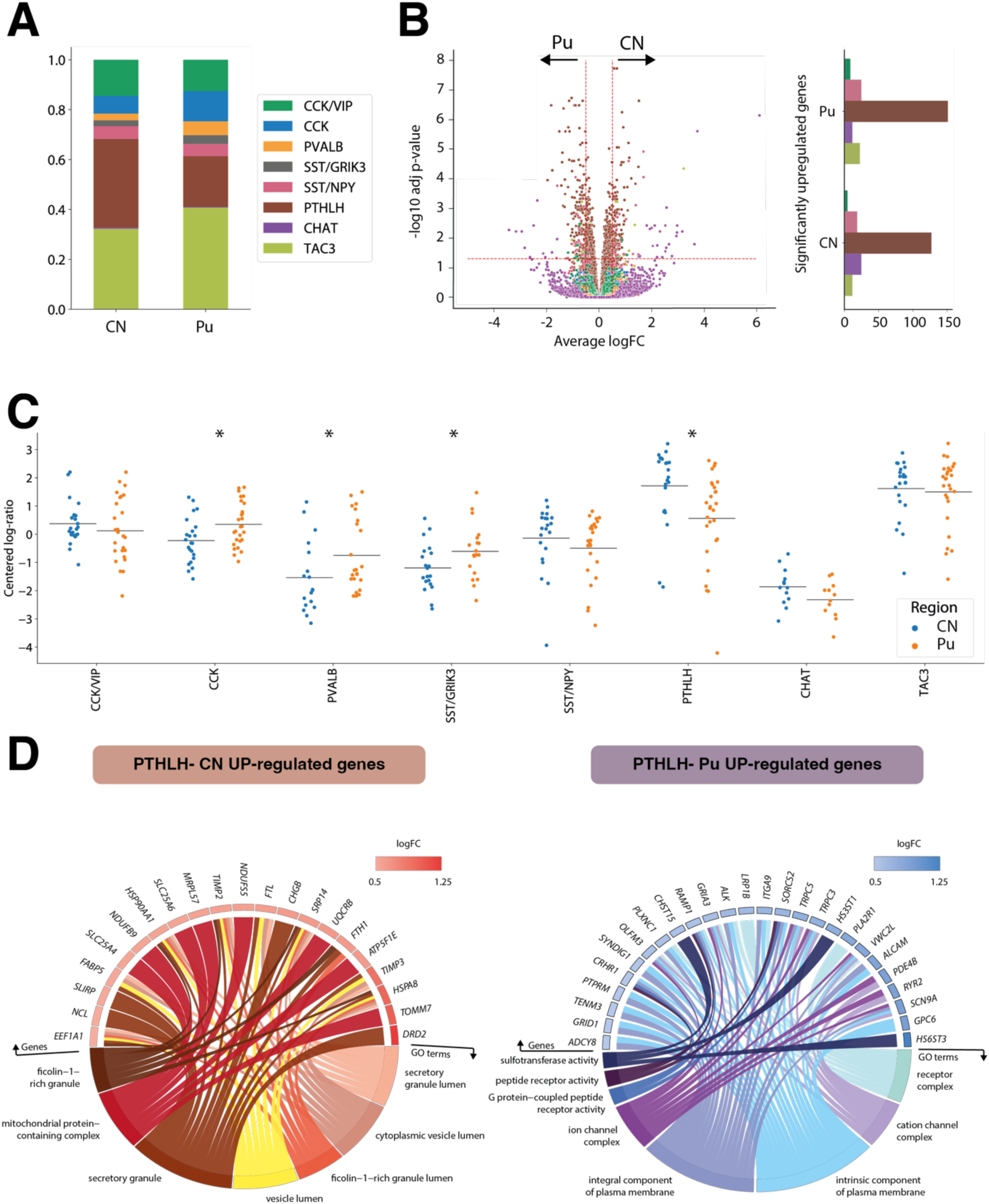
Striatal interneuron classes exhibit region-specific differences. **A.** Barplot illustrating the different proportions of interneuron subclasses in caudate and putamen. **B.** (left) Volcano plot showing differentially expressed genes (DEGs) for each subclass. DEGs with an adjusted p-value < 0.05 and an average logFC greater than 0.5 were selected. (right) Number of significantly upregulated genes per interneuron class on each region. **C.** Scatter dot plot representing the compositional analysis estimated by centered log-ratio method with significant compositional differences between caudate and putamen in CCK, PTHLH, PVALB, and SST/GRIK3 interneurons. **D.** GO-term enrichment analysis of up-regulated genes in PTHLH subpopulation in caudate and putamen. GO circle plot illustrating enriched terms (adjusted p-value < 0.05) with their respective enriched genes along with the logFC of these genes. * P < 0.05.

To contextualize the changes in expression of the PTHLH class neurons across the two striatal regions, we conducted a gene-set enrichment analysis using the genes significantly upregulated in either region (Figure 3D). The 126 genes upregulated in the CN were enriched in GO-terms associated with secretory vesicles and their transport, whereas the 150 genes with significantly higher expression in the Pu were enriched in GO-terms associated with receptor complexes, ion channel complexes, plasma membrane components, and G-protein-coupled receptor activity (GPCR). This signal transduction via GPCRs relies upon the production of cAMP and other signaling cascades^30^. KEGG pathways showed that the differentially upregulated genes in Pu are related to the cAMP signaling pathway, including genes such as *ADCY8* (GPCR), *CRHR1*, *GRIA3*, *GRIN3A*, *PDE4B*, *PLCE1*, and *RYR2*. Thus, these data suggest that there is an over-expression of AMPA and NMDA receptor subunits (related to Ca2+ and Na+ flux) in Pu compared to CN via cAMP/GPCRs activation, which could lead to enhanced long-term potentiation and higher synaptic plasticity in Pu vs. CN and explain why CN and Pu inputs and functionalities are not equivalent^31^.

### PTHLH and TAC3 subclasses exhibit changes along continuous transcriptomic profiles

Our initial cluster analysis allowed us to detect fourteen different interneuron subclasses, each characterized by the expression of a unique combination of marker genes. However, previous studies have shown that striatal interneurons display gene expression gradients^20,21,22^ and, therefore, their diversity may not be captured by oversimplistic binary classifications. To investigate if this phenomenon was observable in our dataset, we conducted a factor analysis within the largest subclasses (PTHLH and TAC3) on each striatal region. In both subclasses and regions, the factor analysis revealed coordinated gradual changes of sets of genes (Figure 4A to D, left and middle panels). The genes with the largest weights on the factor describing the differences within each population were different across regions (Figure 4A to D, right panel, Supplementary table 4), although there were some commonalities; for example, large changes in *SLIT1*, *CNTN5*, and *TAFA2* expression were observed in the PTHLH neurons in both Pu and CN. Similarly, *KCNIP4*, *ASIC2*, and *MARCH1* were responsible for some of the largest variations across the TAC3 neurons in both striatal regions. Notably, some of the genes with the largest contributions to the intra-subclass variance (*KCNIP4*, *ASIC2*, *RYR2*) are ion channel subunits, suggesting the possible existence of different electrophysiological phenotypes within the same transcriptomic subclass. To gain a better understanding of the differences revealed by the factor analysis, we conducted a gene set enrichment analysis on the genes with the largest contributions to the factor on each case (Supplementary Figures 3 and 4).

**Figure 4.**
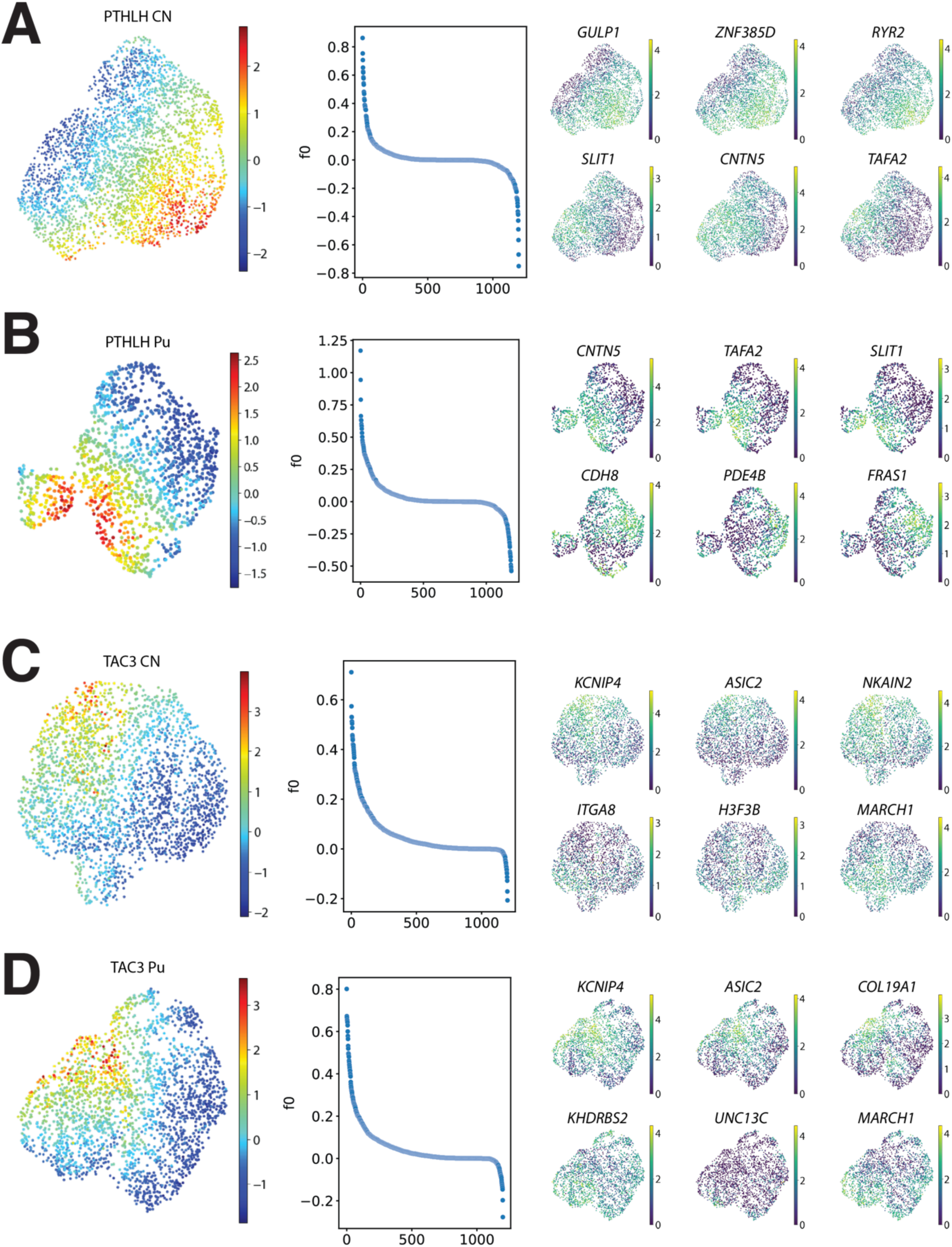
Factor analysis within interneuron subtypes. **A.** (left) UMAP projection of the PTHLH subclass from the CN, colored by the value of the factor obtained by running factor analysis. (middle) Factor weights associated with each gene. (right) UMAP projection of the PTHLH subclass interneurons colored by the expression level of (top row) the genes with the top three (bottom row) and bottom three weights on the factor. **B, C and D.** (left) Factor values, (middle) weights distributions and (right) expression of genes with largest contributions to the factor obtained for the PTHLH subclass from the Pu, the TAC3 subclass from the CN, and the TAC3 subclass from the Pu, respectively.

TAC3 genes associated with the gradient driven by *KCNIP4* and *ASIC2* expression levels showed similar enriched terms in both CN and Pu (Supplementary Figure 4). Relevant terms were mostly related to the regulation of synapse formation or activity and cell adhesion. PTHLH genes related to the gradient driven by *GULP1*, *ZNF385D*, and *RYR2* expression levels in the CN (*CDH8*, *PDE4B*, and *FRAS1* in Pu) displayed terms linked to cell adhesion and channel complexes, specifically Ca^2+^ channels (Supplementary Figure 3). However, results for genes from the Pu gradient defined by *CNTN5*, *TAFA2*, and *SLIT1* showed biological specificity: GO-term enrichment analysis uncovered functionalities in synapse organization and ion transporter activity via ion channels that appear to be specific to the Pu.

### Interneuron taxonomy is maintained across functionally relevant genes

To understand the potential functional implications of our taxonomy, we investigated the differences existing between the established subclasses across two separate sets of genes highly relevant to neuronal function. First, we restricted our dataset to the genes corresponding to dopamine, GABA, acetylcholine, and glutamate receptors. The UMAP projection of our data on this set of genes shows a clear separation between the different subclasses (Figure 5A, left) and the differential expression analysis revealed unique neurotransmitter-receptor expression patterns (Supplementary Figure 5, Supplementary table 5). Markers fitting this pattern noteworthy were: *GRIN3A* and *GRM5* in CCK interneuron class; *GRM7*, *CHRM3*, *GABRA1*, and *CHRNA2* in CCK/VIP; *GRIN2C* in PVALB; *GRIK3*, *GRIK1*, *GRM1*, and *GRIK1* in SST/GRIK3; *GRIP1* and *GRIA4* in PTHLH; *TRPC3* in CHAT; and *GRM8, GRID1, CHRM2*, and *CHRNA7* in TAC3 interneurons.

**Figure 5.**
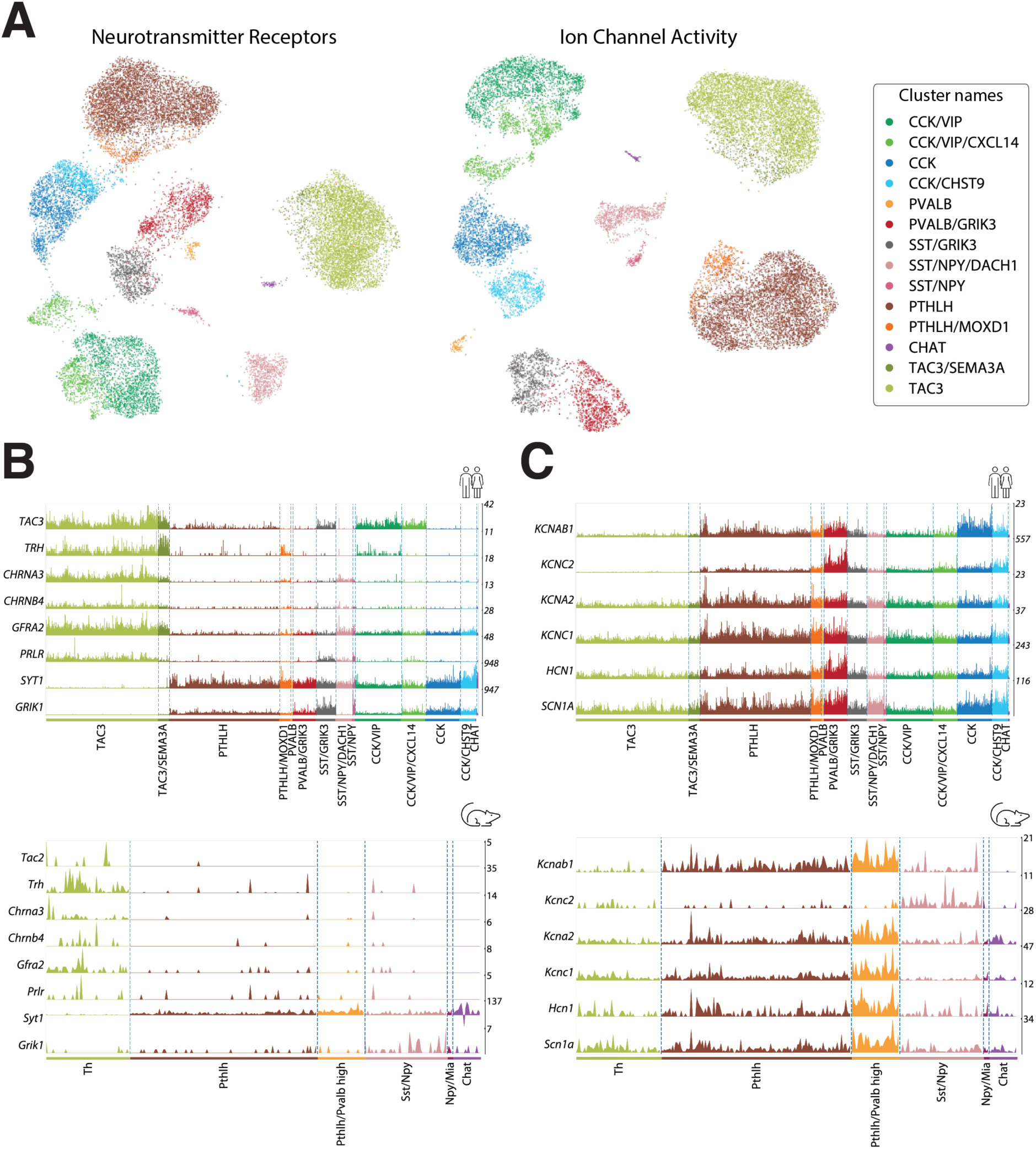
Comparison of striatal interneuron subclasses between mouse and human. **A.** (left) UMAP projection of the interneuron nuclei using expression data restricted to neurotransmitter receptor genes. (right) UMAP projection of the same data restricted to the genes annotated with the molecular function “ion channel activity” (GO:0005216). Colored based on the interneuron classification established before. **B.** (top) Expression values of genes suggesting parallelisms between the TAC3 subclass in the present human dataset and (bottom) the Th interneurons in the mouse striatum described by Muñoz-Manchado et al.^2^. **C.** (top) Expression values of genes related to the fast-spiking phenotype in this human striatal dataset vs. (bottom) the striatal mouse dataset from Muñoz-Manchado et al.^2^

We conducted a similar analysis using all the genes under the GO-term “ion channel activity” (GO:0005216). The UMAP projection of the data using only ion channels retained the separation between subclasses (Figure 5A, right), whereas the differential expression analysis rendered unique transcriptomic patterns. Relevant markers related to these patterns were: *KCNIP1*, *CACNA2D1*, and *KCNH5* in CCK; *GLRA2* and *KCNT2* in CCK/VIP; *KCNMB4* and *RYR1* in PVALB; *GRID2* in SST/GRIK3; *KCNMA1*, *GLRA3*, and *ITPR2* in PTHLH; *TRPC3* and *KCNG3* in CHAT; and *GRID1*, *SCN7A*, and *CACNA2D3* in TAC3 interneurons (Supplementary Figure 5, Supplementary table 5).

A closer inspection of this functional analysis revealed a strong parallelism in key feature genes between the TAC3 population, recently described as a primate-specific class^27^ and that we have thoroughly characterized in the present work, and the mouse interneuron Th cell class^20,14^. Indeed, we found notable expression similarities when comparing the TAC3 human class presented here with the Th mouse class from dataset A in Ref.^20^. Both classes share the expression of the Tachykinin precursor—*TAC3* in human and the homologous gene Tac2 in mouse—and the *TRH* (Thyrotropin Releasing Hormone)—recently described as a marker for the mouse Th population^20^. Additionally, they share the cholinergic nicotinic receptor subunits *CHRNA3/Chrna3* and *CHRNB4/Chrnb4*, the GDNF receptor *GFRA2/Gfra2*, and the prolactin receptor *PRLR/Prlr*. Interestingly, they also match in their negative expression patterns such as the absence of expression of both Synaptotagmin 1 *SYT1/Syt1* and the glutamatergic receptor *GRIK1/Grik1,* which are remarkably highly expressed in the rest of interneuron populations in both human and mouse (Figure 5B). Of note, in agreement with another study in the human striatum^27^, we hardly detect *TH* expression in the human interneuron populations, but we found it in MSNs (data not shown).

To determine the drive of the expression of ion channels in these interneuron subclasses, we focused on the typical genes of a Fast Spiking (FS) profile characteristic of high PVALB-expressing cells (Figure 5C). This analysis showed significant expression of FS genes^32,33–35^ such as *KCNAB1* (Kvb1.3), *KCNC2* (Kv3.2), *KCNA2* (Kv1.2), *KCNC1* (Kv3.1), *HCN1,* and *SCN1A* (Nav1.1) in PTHLH and PVALB cells, with a substantial overexpression in the latter (Figure 5C). In the mouse Pthlh population, the FS profile was found to correlate positively with the *Pvalb* expression level following a continuous gradient pattern Ref^20^. Interestingly, in the human striatum we also found a defined PVALB class of interneurons with high expression of the genes involved in FS profile.

### Interneuron taxonomy is consistent across published human striatal snRNAseq datasets

To further validate our findings and our classification of interneuron subclasses in the human striatum, we integrated our labeled data with four other datasets from three sources^27–29^ using scVI^36^. These datasets included CN^27,29^, Pu^29^, and Nucleus Accumbens^28^ human samples (Supplementary table 6). We filtered and normalized the raw counts from these four datasets and selected the interneuron nuclei based on the same markers as in our own dataset (see Methods). This resulted in a total of 8,090 additional interneuron nuclei added to our 19,339. We reduced the ensemble of the 5 datasets to the 12,986 overlapping genes, of which we selected the top 1,200 most variable to build an integrated model using scVI^36^.

Remarkably, the UMAP projection of the integrated data revealed extensive overlap between nuclei from different datasets, indicating that the the scVI model compensated possible batch-specific differences (Figure 6A). Clustering the integrated data resulted in 16 groups (Figure 6B), of which 15 had a clear correspondence to our original interneuron subclass labels (Figure 6C). Only cluster #12 contained less than 1% of nuclei from our dataset and could not be readily matched to any of the described subclasses. This cluster consisted of 550 nuclei (2% of the total), 83.6% of which belonged to the DropSeq dataset from Krienen et al.’s study^27^. To ensure that cells clustering together actually shared the same transcriptomic profile, we examined the expression of the subclass marker genes on each of the public datasets (Figure 6D, Supplementary figure 6). As the figure shows, cells within the same cluster have similar expression patterns, which in turn correspond to one of the subclasses established in our taxonomy. In the case of cluster #12, we observed that it did express the markers corresponding to our TAC3 subclass. Although there was a strong bias by dataset in some of the clusters, all of them contained nuclei from at least two different datasets, supporting generalizability (Figure 6E).

**Figure 6.**
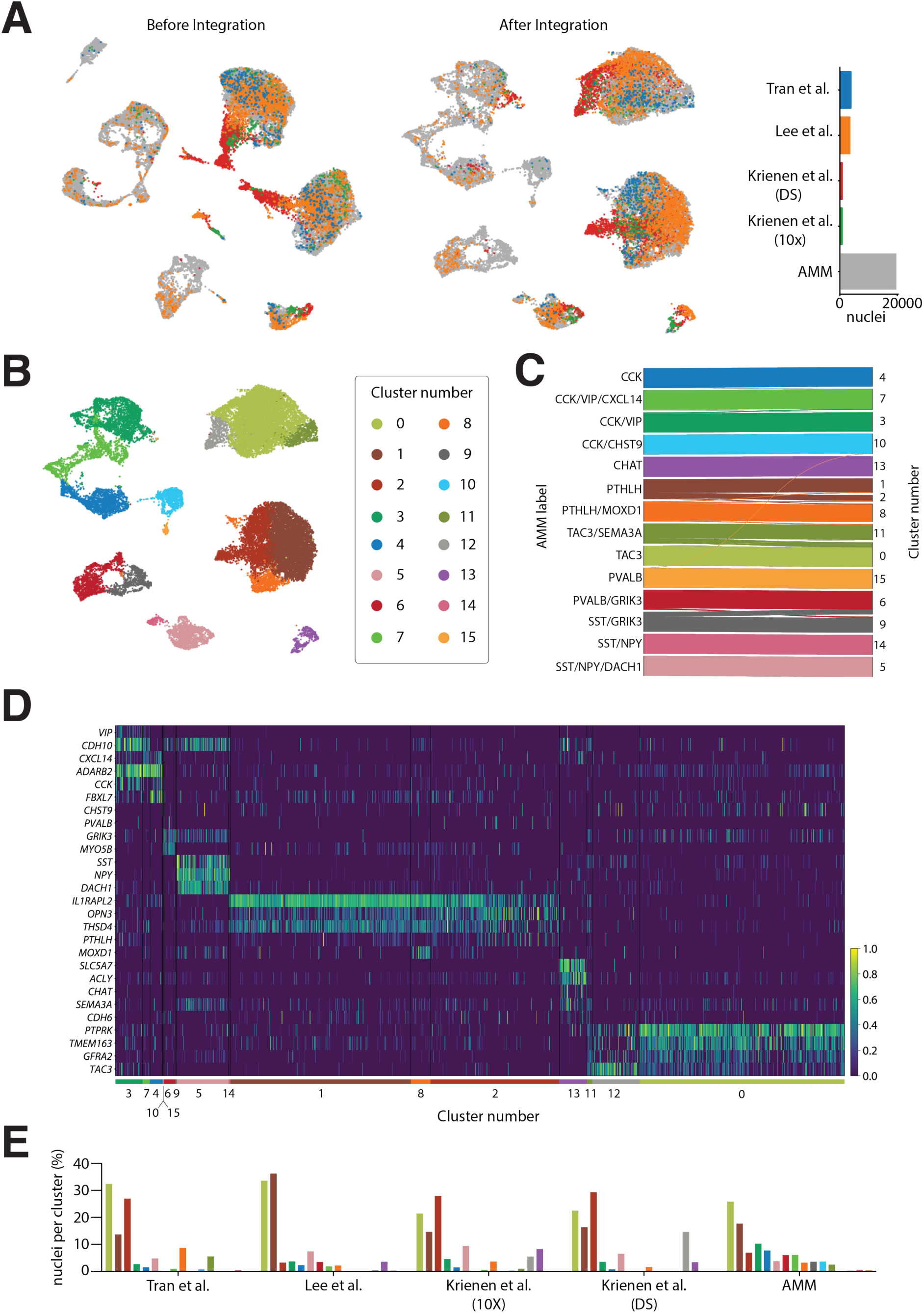
Interneuron taxonomy is consistent across multiple human striatal snRNA-seq datasets. **A.** (left) UMAP projection of interneuron nuclei from five different datasets before and (middle) after integration with scVI. (right) Barplot indicating the total number of nuclei from each dataset. **B.** UMAP projection of the integrated data colored by cluster. **C.** Shankey diagram relating the labels of the nuclei in the AMM dataset to the clusters obtained on the integrated data. Only assignments with more than 1% of the cells of each subclass are shown. **D.** Normalized expression of interneuron subclass marker genes on the integrated public datasets (excluding our own). **E.** In each dataset (*x* axis), each bar represents the percentage of all interneuron nuclei detected in that dataset (*y* axis) that belongs to a specific cluster color-coded as in C.

## Discussion

The neuronal communication in the striatum, a hub for motor and cognitive information, is modulated by the interneurons. Characterizing the diversity and abundance of these locally-projecting neurons is key to understand the proper functionality of this brain structure and here we produced the largest snRNA-seq dataset in number of both nuclei isolated and human samples analyzed to date in order to profile the interneuron diversity of the human dorsal striatum (CN and Pu). We leveraged this dataset to perform a deep molecular characterization of the fourteen interneuron subclasses identified, provide a full set of marker genes for each one, and delineate functional aspects such as synapses-related machinery for different classes, differences between CN and Pu, inner gradient structure of gene expression levels, and relevant pathways related to the main genetic differences.

### Interneuron diversity in the human striatum is higher than expected

The interneuron diversity of the mammalian striatum has received little attention until recently, especially if we compared it with other brain regions such as the cortex where numerous investigations have been carried out^37^. Classically, striatal interneurons have been identified according to five main markers: calretinin (CR or CALB2), Tyrosine Hydroxylase (TH), Parvalbumin (PVALB), Choline acetyltransferase (CHAT), and Neuropeptide Y (NPY)^38,39,24,40^.

With the development of snRNA-seq, many studies have contributed to elucidate the cellular composition of different brain areas including the striatum, especially in the mouse brain. This technology offers a full genetic delineation to characterize molecular cell identities. Only a few published snRNA-seq datasets contain information regarding the human striatum, but they included low number of interneurons and/or focused on other cell types, thus precluding the establishment of a comprehensive taxonomy, as suggested by their lack of agreement^27–29^. In this work, we have sampled nearly half a million CN and Pu nuclei, obtaining almost 20,000 high quality interneuron nuclei after a strict quality control and classification process, a number than enabled us to establish a new taxonomy of human striatal interneurons.

We found eight main classes that we named after one or two of their main molecular markers. Two of these eight main classes express *CCK* and represent almost one third of the total interneuron population. These two *CCK*+ populations were split into two subclasses: CCK/VIP and CCK/VIP/CXCL14 for the ones expressing *VIP* and *CCK,* and CCK and CCK/CHST9 for the ones that do not, respectively. Of note, they all share the expression of *ADARB2*, which separates them from the rest of interneurons and is a marker used to designate development origin from the caudal ganglionic eminence (CGE) of cortical interneurons^41^. Pertinent investigations would be needed to confirm whether this is also the case in striatum, since cortex and striatum seem to present differences in the combinatorial markers of developmental origin^42,43^. CCK and CCK/VIP populations were first reported as striatal interneuron cell class in a scRNA-seq study of the mouse striatum enriched for interneurons^20^. Adarb2/Cck and Adarb2/Vip expressing interneurons have also been described in the striatum of the marmoset and the mouse in a cross-species study^27^. Remarkably, we find an important increase of *CCK*-expressing cells in human vs. mouse dorsal striatum, which might indicate its greater involvement in highly complex computational processes for motor and cognitive functions in the human. Little is known about the role of *CCK*-expressing cells in the central nervous system^44^, although *CCK* is widely used as an interneuron marker in cortical areas. Further investigations would be needed to decipher the role of one of the most abundant interneuron classes in the human striatum.

The hierarchical clustering of subclasses in our taxonomy places the *CCK*-expressing cells in the same main branch as a group of cells expressing the glutamatergic receptor subunit *GRIK3*. These can be divided in three main classes: PVALB, SST/GRIK3, and SST/NPY. The PVALB class further splits into a small population that does not express *GRIK3* (the only ones) and a larger PVALB/GRIK3 population, which represents most cells with a high expression of *PVALB*. Intriguingly, SST/GRIK3 cells are transcriptomically more similar to the *PVALB*+ populations than to the other *SST* expressing cells, which also express *NPY*. The SST/NPY interneurons, one of the classical groups, can be divided by the presence or absence of *DACH1*, which is highly relevant during human neurodevelopment. In the striatum *DACH1* has been described as co-expressed with *SST* as well as several MSN markers^45^. Accordingly, we also observed *DACH1* expression in human striatal MSNs in adulthood (data not shown).

In the other branch of our classification, we find the two populations representing the most abundant interneuron classes: PTHLH and TAC3, together with the well-described and scarce cholinergic cells (CHAT), which have already been thoroughly characterized in the literature^3,25,46,47^. Both PTHLH and TAC3 populations further split into two subclasses of unequal proportions: PTHLH and PTHLH/MOXD1, and TAC3 and TAC3/SEMA3A, respectively.

A *PTHLH*+ population was first described in the mouse striatum^20^, where it was characterized as a group of cells that expressed Pvalb in a gradient manner that correlated with their electrophysiological properties, spatial distribution, morphology, and long-range inputs^23^. In the human striatum this population is characterized by the expression of *OPN3*, *IL1RAPL2*, and *THSD4*, and shows a higher abundance in the CN than in the Pu. A PTHLH subclass shows specific expression for MOXD1—a monooxygenase predicted to be involved in the dopamine catabolic process—suggesting dopaminergic modulation by these cells. Similarly, to the mouse striatum, we find *PVALB* expression in the PTHLH population. This *PTHLH*+/*PVALB*+ population has also been confirmed by others in human striatum and other species such as the marmoset and the mouse amygdala^27^, https://www.biorxiv.org/content/10.1101/2022.10.25.513733v1. Interestingly, we have found a specific and less abundant class that expresses *PVALB* (at a significantly higher level than the PTHLH/PALB cells) but not *PTHLH*. This finding was validated with FISH and differs from the mouse striatum, where all Pvalb-expressing cells were also Pthlh+. These two classes, PTHLH and PVALB, do not appear close in their molecular identities in the human striatum when applying hierarchical analysis; however, when we performed a hypothesis-driven analysis of our data, restricted to relevant genes for neuronal functions such as neurotransmitter receptors or ion channels, these two classes showed a very strong correlation. This suggests that even though their overall molecular identities are far apart, these two classes might share functional roles in the striatal circuit. This observation brings up the recurrent debate on what constitutes a class^48–51^ and, more importantly, indicates that examining specific aspects of cell identity will deliver different pieces of information, such as what other cell(s) they communicate with and what kind of electrical activity they present. With that framework in mind, we also analyzed the most relevant genes for Fast Spiking activity, characteristic of high Pvalb-expressing cells in the mouse striatum. Our data showed that in the human striatum all the Fast Spiking relevant genes were differentially upregulated in both PTHLH and PVALB classes, but substantially higher in the latter.

TAC3 was recently described as a primate-specific striatal population^27^. We did define a population of interneurons with high *TAC3* expression as the TAC3 class. However, this class was best defined by the expression of *PTPRK*, *TMEM163*, and *GFRA2*, since *TAC3* is also expressed by the CCK/VIP class, a co-expression that was also reported by Krienen et al.^27^. Through our functional gene analysis, we observed that TAC3 is characterized by synaptic receptors for glutamate (*GRM8*) and acetylcholine muscarinic (*CHRM2)* and nicotinic (*CHRNA7*, *CHRNB4)* receptors. Among the genes with ion channel activity, we found *GRID1* (glutamate ionotropic receptor), *SCN7A* (sodium voltage-gated channel), and *CACNA2D3* (Calcium Voltage-gated channel). Interestingly, when comparing functional genes in human vs. mouse striatum^20^, we found that the mouse interneuron Th cell class had been previously described to express nicotinic receptors, including those responding to a3b4 (a specific subtype)^47,52,53^. This mouse Th population is characterized by the expression of *Tac2* (homologous of the human *TAC3* gene coding for a tachykinin precursor) and also by the expression of *Trh*, which is one of the best markers for Th cells in mouse. Remarkably, both populations share a specific pattern expression for *PRLR* (prolactin receptor), *GFRA2* (GDNF receptor), *SYT1* (synaptotagmin), and *GRIK1* (Glutamate ionotropic receptor subunit). For the last two genes (*SYT1* and *GRIK1)*, both involved in synaptic function, TH and TAC3 populations present a negative pattern, being the absence of SYT1 and GRIK1 what differentiate them in a remarkably manner from the other interneuron populations in the striatum. Although integration of human and mouse datasets was not technically possible, even despite applying recent tools as LIGER^54^, the aforementioned genes showed a strong parallelism between the mouse Th and the human TAC3 populations. Importantly, in agreement with others^27^, we hardly found *TH*-expressing interneurons, although we did find TH expression in MSNs as shown elsewhere^32,55,56^. This observation points out that TH expression in striatum probably cannot be used as marker for this cell class, at least in humans, and suggests that, from the evolutionary perspective, the absence of TH in the TAC3 population might just indicate a refinement in the circuitry or a loss of unnecessary machinery. Noteworthy, since TH is the limiting enzyme in the synthesis of dopamine and noradrenaline, we also examined our data for genes related to dopamine metabolism and found none (data not shown).

### Inner gradient structure is conserved in the human striatum

Because the discrete partitions of the data might not reveal the entire actual biologically relevant diversity of the striatal interneuron populations^49^, we examined the PTHLH and TAC3 subclasses using factor analysis. With this approach, we did find that there is diversity within the subclasses, as indicated by differences in expression patterns along a continuum rather than by discrete changes in the expression of a set of marker genes. The gradients within each subtype were driven by the same set of genes in both the CN and the Pu. We also studied the gradient structure in the human striatum as previously shown in the mouse striatum for both interneuron and MSNs^20–22^. We found an inner gradient structure for the most abundant classes, TAC3 and PTHLH, which is shared in both CN and Pu, indicating similarities in structure, as it was shown for Pthlh in the mouse striatum Ref20. This gradient structure seems to be characteristic of subcortical structures such as the striatum and may reflect the need of a highly specialized organization to receive input from many different and distant brain areas.

### Differences across regions

The main differences identified between CN and Pu in our study are related to the PTHLH class. Our results indicate that this cell class in the Pu might be involved in long-term potentiation mechanisms, a form of synaptic plasticity that plays a critical role for the proper functionality in the striatum^9,57^. This difference may underscore a potential different vulnerability of Pu vs CN to basal ganglia-related diseases, as already suggested by others^58–60^, which could be used in the design of cell type-targeted therapeutic approaches.

### Robustness of striatal interneuron taxonomy

Besides performing *in situ* tissue validations of the interneuron classes and subclasses, we validated our taxonomy by integrating our data with previously published sn/cRNA-seq datasets. An unbiased clustering of the integrated data resulted in groups of interneurons which could be readily overlapped or at least related to each of the subclasses we describe here and, more relevant, the expression profile of marker genes within each group was consistent across datasets, even in the non-integrated raw data. This indicates that our taxonomy is robust, as even the rarest cell subclasses could be observed in other datasets. Most notably, the classification we introduce was highly compatible with the samples from the nucleus accumbens (ventral striatum) from Tran et al^28^, suggesting that the inhibitory neurons in both ventral and dorsal striatum share a similar diversity spectrum. However, a broader sampling of ventral striatum would be useful to reinforce this observation and to determine if this classification can be extended also to other regions in the basal ganglia.

## Material and Methods

### Human tissue

Postmortem human Pu (N = 28) and CN (N = 25) fresh frozen tissue samples from 28 control donors aged 25 to over 90 years were obtained from three sources, the NIH Neuro Bio Bank (Human Brain and Spinal Fluid Resource Center Los Angeles, CA, USA), the Parkinson’s UK Brain Bank (London, UK) and the Massachusetts Alzheimer Disease Research Center (Charlestown, MA, USA). Sample information can be found in supplementary table 1.

### Tissue dissociation

Isolation of nuclei from fresh frozen tissue was performed as described by the Allen Institute for Brain Science (https://www.protocols.io/view/isolation-of-nuclei-from-adult-human-brain-tissue-eq2lyd1nqlx9/v2) with the following specifications, all steps were performed at 4°C. 100 – 150 mg of tissue was thawed on ice and homogenized in 2 ml of chilled, nuclease-free homogenization buffer (10 mM Tris (pH 8), 250 mM Sucrose, 25 mM KCl, 5 mM MgCl_2_, 0.1 mM DTT, 1x Protease inhibitor cocktail (50x in 100% Ethanol, G6521, Promega), 0.2 U/µl RNasin Plus (N2615, Promega), 0.1% Triton X-100) using a Dounce tissue grinder with loose and tight pestle (20 strokes each, 357538, Wheaton). The nuclei solution was filtered through 70 µm and 30 µm strainers successively, tubes and strainers were washed with an additional homogenisation buffer (final volume 6 ml) before centrifugation for 10 min at 900 rcf. Supernatant was removed leaving 50 µL above the pellet and resuspended in 200 µl homogenization buffer (final volume 250 µL). Then, the suspension was carefully mixed 1:1 with 50% Iodixanol (OptiPrep Density Gradient Medium (D1556, Sigma) in 60 mM Tris (pH 8), 250 mM Sucrose, 150 mM KCl, 30 mM MgCl_2_) and layered carefully on top of 500 µL 29% Iodixanol in a 1.5 ml tube. Samples were spun 20 min at 13,500 rcf and supernatant was removed as much as possible without disrupting the pellet. Pellet was resuspended in 50 µL chilled, nuclease-free blocking buffer (1x PBS, 1 % BSA, 0.2 U/µL RNasin Plus), transferred to a fresh tube and filled up to 500 µL. To enable enrichment of neurons during fluorescent activated cell sorting, 1µl NeuN antibody (1:500, Millimark mouse anti-NeuN PE conjugated, FCMAB317PE, Merck) was added and samples were incubated for 30 min on ice in the dark. After spinning 5 min at 400 rcf, the supernatant was removed leaving ∼50 µL of buffer above the pellets and 500 µL of blocking buffer was added to resuspend before filtering through a 20 µm filter into FACS tubes and adding 1 µL of DAPI (0.1 mg/mL, D3571, Invitrogen).

### Fluorescent-activated nuclei sorting

Nuclei suspension was protected from light and sorted in a flow cytometer (DB FACSAria Fusion or BD FACSAria III) at 4°C. Gating was performed based on DAPI and phycoerythrin signal into two tubes containing 50 µL blocking buffer (NeuN+ and NeuN-population) until 200,000 nuclei per population were reached. Sorted populations were centrifuged 4 min at 400 rcf and supernatant was removed, leaving approximately 30 µL to resuspend the pellet, samples were kept on ice.

### single-nucleus RNA sequencing library preparation

Library preparation from sorted nuclei suspension was done using the Chromium Next GEM Single Cell 3’ Reagent Kit v3.1 (PN-1000268, 10x Genomics). Each nuclei population was counted manually, and the concentration was adjusted to a range between 200 and 1,700 nuclei/µL. Following the manufacturer’s protocol (CG000204 Rev D, 10x Genomics), RT mix was added to the nuclei suspension and samples were either loaded for each population on separate lanes (target nucleus recovery 5,000) or population were mixed (70% NeuN+ and 30% NeuN, target nucleus recovery 5,000 or 7,000) before loading on one lane of the Chromium Next GEM Chip G (PN-1000120, 10x Genomics). Downstream cDNA synthesis and library preparation followed the manufacturer’s instructions using the Single Index Kit T Set A (PN-1000213, 10x Genomics). Required quality control steps and quantification measurements within this protocol were performed using the Agilent High Sensitivity DNA Kit (5067-4626, Agilent Technologies) and the KAPA Library Quantification Kit (2700098952, Roche).

### Illumina sequencing

Pools were prepared by combining up to 19 (target nucleus recovery 5,000) or 16 (target nucleus recovery 7,000) samples and sequencing was performed on a NovaSeq S6000 using a S4-200 (v1.5) flowcells with 8 lanes and a 28-8-0-91 read set up. The sequencing was performed at the National Genomics Infrastructure (Stockholm, Sweden).

### Tissue preparation for histology

Human tissue blocks (N = 6) were stored at −80 °C and transferred to the cryostat (CryoStar NX70, Thermo Scientific) on dry ice. Samples were mounted on the specimen holder using Tissue Tek O.C.T. Compound (4583, Sakura) and acclimated to −20°C in the cryostat chamber for 5 minutes. 10 µm sections were cut and captured on Super-Frost Plus microscope slides (631-0108, VWR) at room temperature. Slides were air dried at room temperature for a few minutes and stored for 1h at −20°C before transferring them back to −80°C for long term storage.

### RNAscope high sensitivity *in-situ* hybridization

High sensitivity *in situ* hybridization using the RNAscope Multiplex Fluorescent Reagent Kit v2 (323110, Advanced Cell Diagnostics) was performed on putamen sections of six subjects to detect single mRNA molecules. Experiments were performed according to the RNAscope Multiplex Fluorescent Reagent Kit v2 protocol (UM 323100, Advanced Cell Diagnostics) for the following genes: *DACH1* (412041), *NPY* (416671-C2), *PTHLH* (452931), *PVALB* (422181-C2), *SST* (310591-C3). In brief, slides were dried at room temperature (RT) for 5–10 min before incubation in 4% PFA for 25 min at 4°C. Slides were washed twice in 1x PBS and dehydrated in 50%, 70% and 2x 100% ethanol for 5 min each at RT. After drying the slides for 5 min, a hydrophobic barrier was drawn around each section prior to incubation in hydrogen peroxide for 10 min at RT. For antigen accessibility, slides were treated with Protease IV for 20 min at RT after a brief wash in 1x PBS. Slides were washed twice for 3 min again, before probes were incubated. C2 and C3 probes were diluted in C1 probes at a 1:50 ratio and incubated on the slides for 2h at 40°C. Slides were then incubated with amplification mix 1-3 followed by a combination of HRP reagent, fluorescent dye and HRP blocker specific for each probe channel in accordance to the manufacturer’s recommendations. Probes were detected with Opal 520 (FP1487001, Akoya Biosciences), Opal 570 (FP1488001, Akoya Biosciences) and Opal 650 (FP1496001, Akoya Biosciences). Next, slides were incubated with TrueBlack (23007, Biotium) after a wash in 70% ethanol for 30 sec at RT to quench the autofluorescence due to the accumulation of lipofuscin or other protein aggregates. Prior to mounting with Fluoromount-G (0100-01, SouthernBiotech) the slices, DAPI was added to label the nuclei. A one-day protocol has been used in all experiments to preserve the quality of the slices.

### Image acquisition

Confocal imaging was performed on a Zeiss LSM800-Airy with Zen software (2.6). 2-3 non-overlapping areas with a size of 8 x 8 tiles were selected per tissue section and images were acquired using a 20x air/dry objective. Final images were stitched using the according feature of the Zen software.

### Image analysis

Quantitative image analysis was performed using the QuPath software (version 0.3.2)^61^ with the following workflow: (1) Definition of region of interest (ROI) on each image across all visible nuclei (using DAPI stain) but excluding artifacts and high fluorescent vessels, based on size and intensity of the signal. (2) Cell and subcellular detection tools of QuPath were adjusted for each staining (supplementary table 7) and applied within each ROI. (3) An object classifier was trained for each marker and subject individually by manual labeling of positive cells. Cells were considered positive when either a clear fluorescent signal was visible throughout the approximated cell body (*NPY*, *SST*) or a distinct puncta signal was evident with no overlapping signal in other fluorescent channels (*DACH1*, *PTHLH*, *PVALB*). The purpose of using object classifiers instead of counting subcellular spots directly was to improve the distinction between truly positive cells and cells with autofluorescent signal due to lipofuscin. (4) All relevant object classifiers for a specific staining and subject were combined to a composite classifier and applied to all ROI of the respective subject. (5) The resulting list of cells and their assigned markers per ROI were exported and evaluated for each subject within a staining. Among the group of positive cells minimum cut off values concerning the number of subcellular spots were defined for each marker (supplementary table 7) and applied manually. Additionally, unexpected marker combinations or numbers of spots were checked manually on the image and corrected if necessary.

### snRNA-seq data pre-processing

The raw data was processed into count matrices by using CellRanger (v.3.0.0) (10X genomics) to align the sequencing data to the hg38 genome (GRCh38.p5 (NCBI:GCA_000001405.20), accounting for both intronic and exonic sequences.

To detect possible doublets, we applied Scrublet^62^ to each individual sample 100 times with automated threshold value detection and default parameters. Nuclei labeled as doublets more than 10 times were discarded.

Based on the distribution of UMIs and unique genes detected per nucleus, cells with less than 500 UMIs or 1,200 genes were discarded. Cells with more than 250,000 UMIs, over 15,000 genes or more than 10% mitochondrial content were also excluded.

Using the nuclei which passed the initial QC thresholds, we modeled the relationship between number of unique genes and UMIs in the logarithmic scale as a second-degree polynomial function. Nuclei with extreme deviations from the polynomial fit (a difference over 2,000 between log (n genes) and the value predicted by the fit for a given UMI count) were considered outliers and excluded from the rest of the analysis.

Cells expressing high levels of marker genes for multiple cell types simultaneously were also discarded. To do so, we computed a cell-type score for each cell subtype (Oligodendrocytes, Microglia, OPCs, Neurons, Astrocytes, Vascular cells) for each nucleus. This score was the mean expression of the canonical markers for each type. Then, we computed the distribution of the scores on the whole dataset, observing bimodal distributions in all cases. We modeled the distributions as mixtures of two Gaussians and set a threshold on the mean of the lowest distribution plus four times its standard deviation. Nuclei with a score above the threshold were considered of a given cell type. Nuclei with scores above the threshold for more than one type were considered doublets and excluded from the rest of the analysis.

To remove possible contamination from the claustrum or the amygdala, we removed cells expressing regional markers obtained from the Allen Brain atlas^63^: *NEUROD2*, *TMEM155*, *CARTPT*, *SLC17A7*.

The number of nuclei excluded at each step of this process is detailed in Supplementary figure 1.

### Interneuron detection

The count matrices were analyzed using Scanpy^64^ to cluster and label them in order to select for interneurons. Briefly, we performed principal component analysis and computed the neighborhood graph on the first 30 principal components (PCs). The data was then clustered using the Louvain algorithm^65^ with a resolution of 0.2 and the clusters were labeled as either glia or neurons based on the expression of canonical markers:

Astrocytes – *AQP4*, *ADGRV1*

Microglia – *CSF1R*, *FYB1*

Oligodendrocytes – *MBP*, *MOG*, *MAG*

Oligodendrocyte precursor cells – *PTPRZ1*, *PDGFRA*, *VCAN*

Vascular cells – *EBF1*, *ABCB1*, *ABCA9*

Neurons – *MEG3*

The neurons were filtered again based on their distribution of UMIs and genes. Nuclei labeled as neurons with less than 5,000 UMIs, less than 3,000 genes or more than 12,000 genes were discarded, resulting in a total of 181,434 high quality neuronal nuclei.

The neurons were re-clustered after removing sex-linked mitochondrial and riboprotein genes, projecting them onto their first 30 PCs computed on their 1,500 most variable genes. The clusters expressing the inhibitory markers *GAD1* and/or GAD2 and not expressing MSN (*PPP1R1B*, *DRD1*, *DRD2*, *MEIS2*) or excitatory markers (*RORB*) were labeled as interneurons.

### Interneuron classification

Nuclei labeled as interneurons were projected onto the first 20 PCs calculated on their 1500 most variable genes and re-clustered using the louvain algorithm. The function *rank_genes_groups* from Scanpy^64^ was used to perform a differential expression analysis between the clusters through a Wilcoxon rank-sum test. Marker genes were selected manually from the top ranked genes to characterize and name each of the interneuron clusters as a different interneuron subclass.

The interneuron subtypes were merged into broader classes based on their correlation. All subtypes with a mean Pearson correlation coefficient higher than 0.49 to each other were joined into a broader class defined by common marker genes.

The dendrograms were computed using the average Pearson correlation coefficient between groups across all genes.

### Compositional analysis

The differences in composition between the CN and the Pu were examined through the centered-log ratio (CLR) values for each interneuron class on each of the regions. This measure is defined as

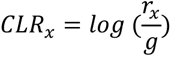

Where r_x_ is the fraction of interneurons of a given class and g is the geometric mean of the fractions of each of the classes.

The distribution of CLRs for the same interneuron class were compared across regions using a non-parametric Wilcoxon test.

### Differential expression analysis (DEA)

Regional changes in gene expression were studied using a pseudo-bulk approach in which the nuclei were aggregated by region and sample. The Libra python library^66^ was used to perform the data aggregation and the differential expression analysis, which was done using the edgeR-LRT method^67^.

In all the other cases, differential gene expression was studied at the cell level using the *rank_genes_groups* function from the Scanpy library using a Wilcoxon rank-sum test.

### Over-Representation Analysis (ORA)

Differentially Expressed Genes (DEGs) derived from DEA were used as input into the enrichGO and enrichKEGG functions from the R package clusterProfiler. DEGs with a logFC > 0.5 and p-adjusted value < 0.05 were selected. The first function generates functional Gene Ontology terms related to biological processes, molecular function, and cellular components. The second function analyzes the enriched terms in our gene list based on the KEGG database (DB). This DB is a collection of manually drawn pathway maps representing our knowledge of the molecular interaction, reaction, and relation networks for Metabolism, Genetic Information Processing, Environmental Information Processing, Cellular Processes, Organismal Systems, Human Diseases, and Drug Development. Terms with a p-value < 0.1 were selected and plotted using the GOplot package.

### Factor analysis

The heterogeneity within the PTHLH and TAC3 subclasses was studied using a factor analysis. For each interneuron subclass on each of the two striatal regions, we removed the sex-linked, mitochondrial and riboprotein genes, and then restricted the data to the 1,200 most variable genes. We then applied the *FactorAnalysis* function from the scikit-learn Python library^68^ with a single latent factor to perform a matrix decomposition.

### Data projection on functional gene subsets

To restrict the data to neurostransmitter-receptor genes, we selected genes based on their prefixes: *DRD*-(dopamine); *GABR*-(GABA); *CHRN*-, *CHRM*-(acetylcholine); *GRIA*-, *GRIN*-, *GRIK*-, *GRM*-, *GRID*-, *GRIP*-(glutamine). We added three additional glutamine receptors whose naming did not follow the same pattern: *PEPL1*, *POLR2M*, *GCOM1*. This selection resulted in 93 genes.

To study the genes with ion channel receptors, we restricted our data to the genes listed under the GO-term GO:0005216. This list contained 481 genes names, out of which 431 were found in our data.

In both cases, we obtained the UMAP projection from the neighborhood graph computed on the first 30 PCs and then performed a differential expression analysis using a Wilcoxon rank-sum test.

### Public datasets collection and pre-processing

We collected two single-nuclei RNA-seq datasets of the human striatum from the GEO database^69^, with accession numbers GSE151761^27^ and GSE152058^29^. A third dataset was obtained from a public repository setup by the authors^28^ (https://github.com/LieberInstitute/10xPilot_snRNAseq-human). On Krienen *et al*.’s data, we analyzed separately the 10X and Drop-Seq datasets. On Lee *et al*.’s data we used only the nuclei belonging to control subjects (8 samples). We normalized all the datasets using scran normalization^70^ and applied individual QC filters to remove bad quality nuclei. We then clustered the data and selected the interneuronal populations using the same approach and criteria that we applied to our own data. Notably, on Lee *et al*.’s dataset our selection included a cluster originally labeled as secretory ependymal cells, which expressed both pan-neuronal and interneuronal markers, and we identified as TAC3 interneurons. The total number of cells filtered and selected are detailed in supplementary table 6.

The mouse data from Muñoz-Manchado *et al*.^20^ was obtained from the GEO database (accession number GSE97478). The raw counts were normalized using the *normalize total* function from Scanpy, with a target sum of 10000 per cell. The original labels were retained, and the data was not transformed further.

### Data integration

snRNA-seq dataset from multiple sources were integrated using scVI^36^. First, the data was merged and restricted to the 12986 genes common across datasets. Then the 1200 most variable genes were selected and used to build and train an autoencoder with 1 hidden layer of 128 nodes and a latent space of dimensionality 12 which was trained for 292 epochs. The low-dimensional latent state representation was used to build a neighborhood graph and then cluster the data in the same way as on the PC-projected data from our novel dataset.

## Acknowledgments

The authors thank Patrick Dooley and Tessa Connors from the Massachusetts Alzheimer’s Disease Research Center for their assistance in selecting the donors. The authors acknowledge the Parkinson’s UK Brain Bank (Requestor: A.B.M.-M) and NIH NeuroBioBank (1851-Requestor: Ernest Arenas). The authors acknowledge support from the Spanish Ministry of Science and Innovation (I+D+I; RyC), the Swedish Foundation for Strategic Research, the Swedish Research Council, and the US National Institute on Aging. The authors also want to thank the National Genomics Infrastructure in Stockholm funded by Science for Life Laboratory and SNIC/Uppsala Multidisciplinary Center for Advanced Computational Science for assistance with massively parallel sequencing and access to the UPPMAX computational infrastructure. We also thank the donors and their families who have contributed to this study.

## Author contributions

L.G., L.H. and A.B.M.-M designed the study. L.H., M.D.-S and A.B.M.-M carried out experiments, L.G., J.M.B.-R, L.H. and A.B.M.-M performed data analysis. A.S.-P. and B.T.H provided resources. L.G., L.H. and A.B.M.-M wrote the manuscript with comments and input from all authors.

## Data availability

All raw sequencing data and custom code will be made available upon publication.

