## Supplementary figures for "Interneuron diversity in the human dorsal striatum"

**Supplementary Figure 1**

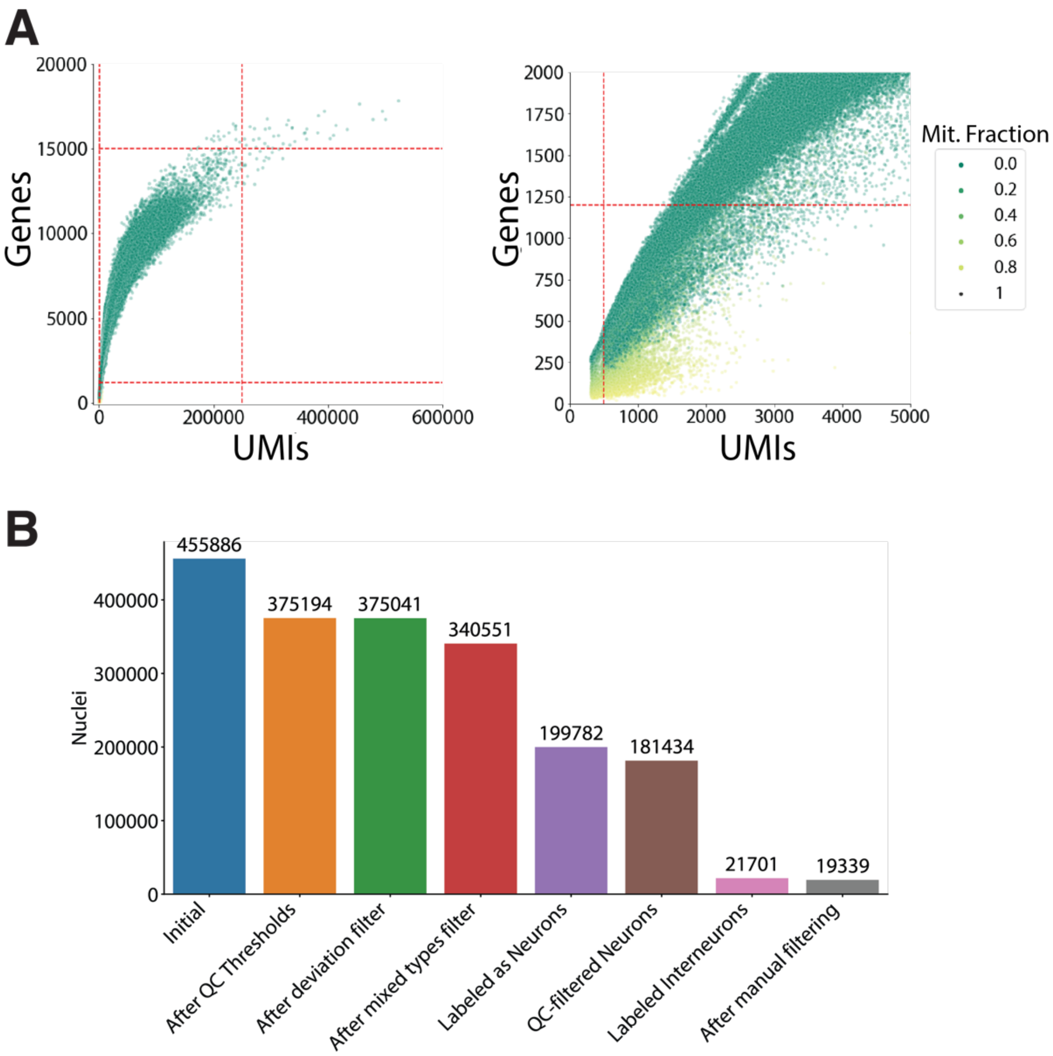


**Supplementary Figure 1. Quality control metrics of the snRNA-seq data. A.** Distribution of the number of genes and counts per nucleus, colored by the fraction of total detected UMIs corresponding to mitochondrial gene products. The values chosen for the initial thresholds are shown in red (see Methods). **B.** Number of nuclei retained at each step of the analysis.

**Supplementary Figure 2**

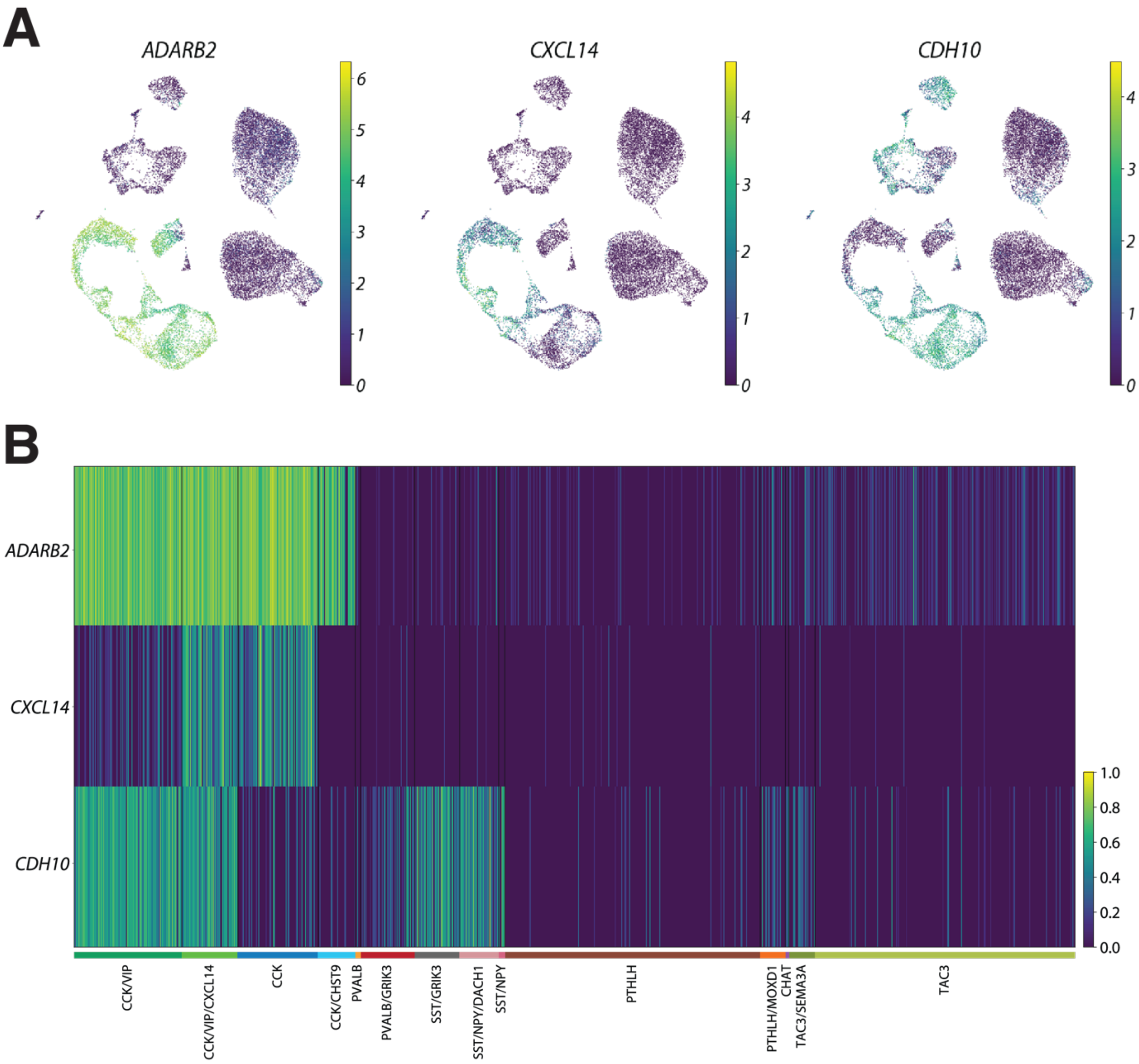


**Supplementary Figure 2. Expression of *CXCL14* and *CDH10* distinguishes *ADARB2*+ interneuron subtypes. A.** UMAP projection of all the interneuron nuclei showing the normalized expression of *ADARB2*, *CXCL14* and *CDH10*. B. Normalized expression of the same genes on the nuclei ordered by interneuron subtypes. As the plot shows, *CXCL14* and *CDH10* could be used to distinguished the different subclasses of *ADARB2*+ interneurons: *CDH10*+/*CXCL14*- (CCK/VIP); *CDH10*+/*CXCL14*+ (CCK/VIP/CXCL14); *CDH10*-/*CXCL14*+ (CCK); *CDH10*-/*CXCL14*- (CCK/CHST9).

**Supplementary Figure 3**

**A** GOterms - PTHLH gradient (CN)

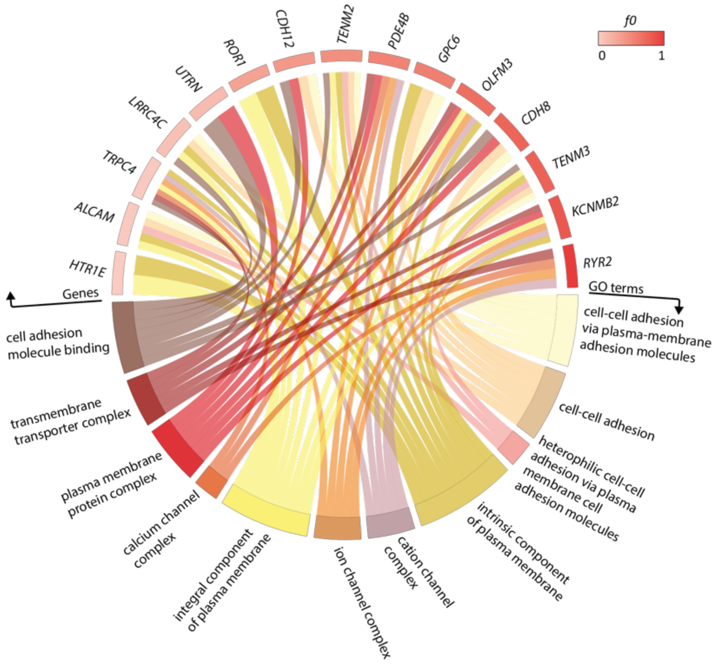


**B** GOterms - PTHLH gradient (Pu)

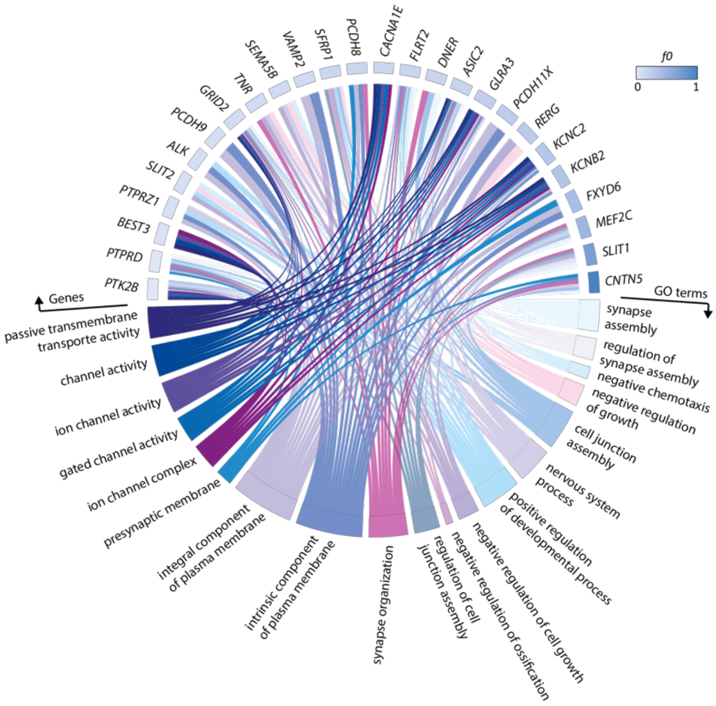


**Supplementary Fig. 3 | GO enrichment analysis of genes correlated positively with the PTHLH vector from the factor analysis. A.** GO circle plot representing enriched terms (adjusted p-value < 0.05) with their respective enriched genes along with the f0 value of these genes. Genes correlated positively with the vector in PTHLH subclass in Caudate (f0 > 0.2) were selected. **B.** Same as A but with genes correlated positively with the vector in PTHLH subclass in putamen. Both sets of genes show similar GO-terms enrichment.

**Supplementary Figure 4**

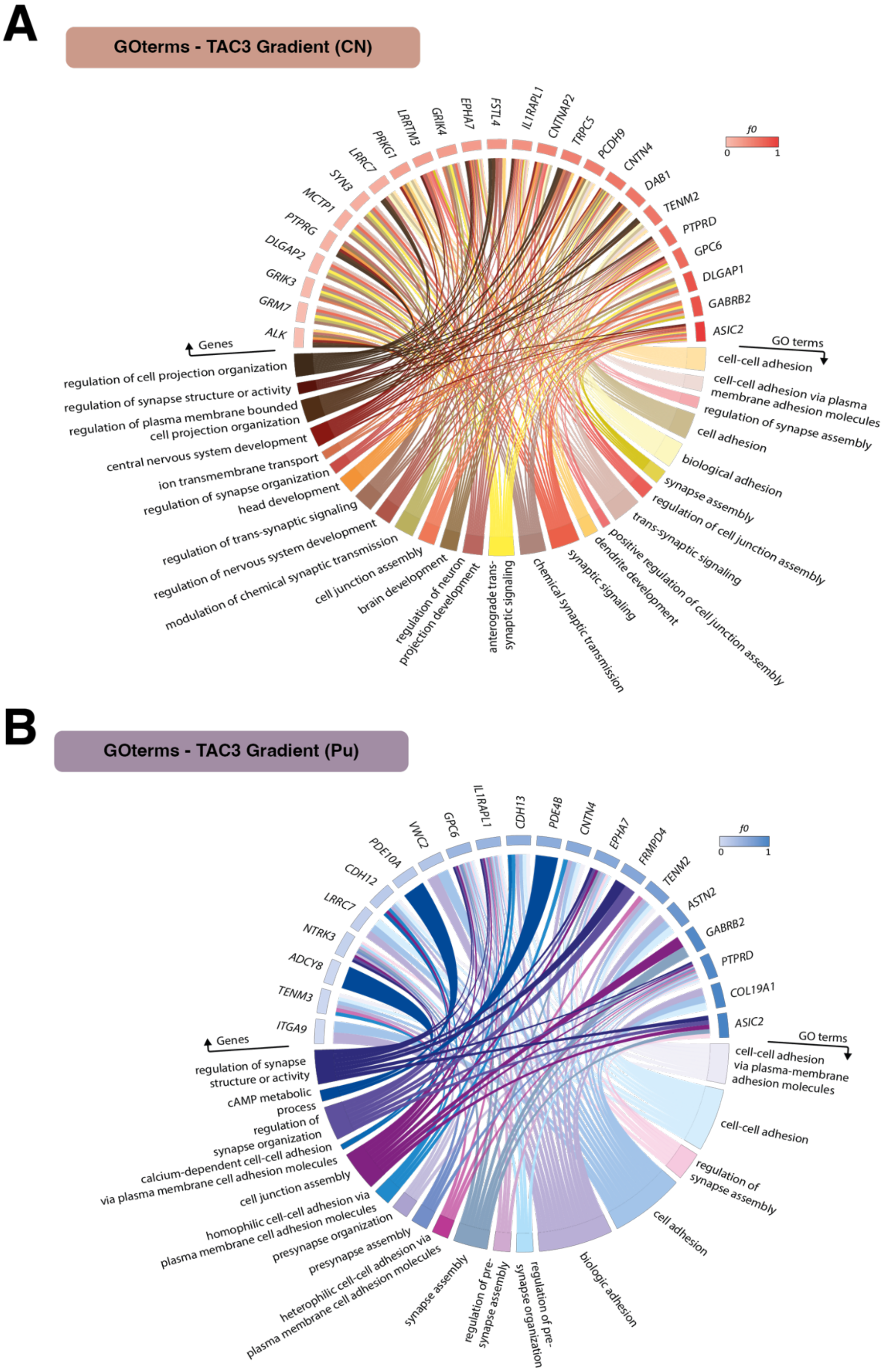


**Supplementary Fig. 4 | GO enrichment analysis of genes correlated positively with the TAC3 vector from the factor analysis. A.** GO circle plot representing enriched terms (adjusted p-value < 0.05) with their respective enriched genes along with the f0 value of these genes. Genes correlated positively with the vector in TAC3 subclass in caudate (f0 > 0.2) were selected. **B.** Same as A but with genes correlated positively with the vector in TAC3 subclass in putamen. Both sets of genes show similar GO-terms enrichment.

**Supplementary Figure 5**

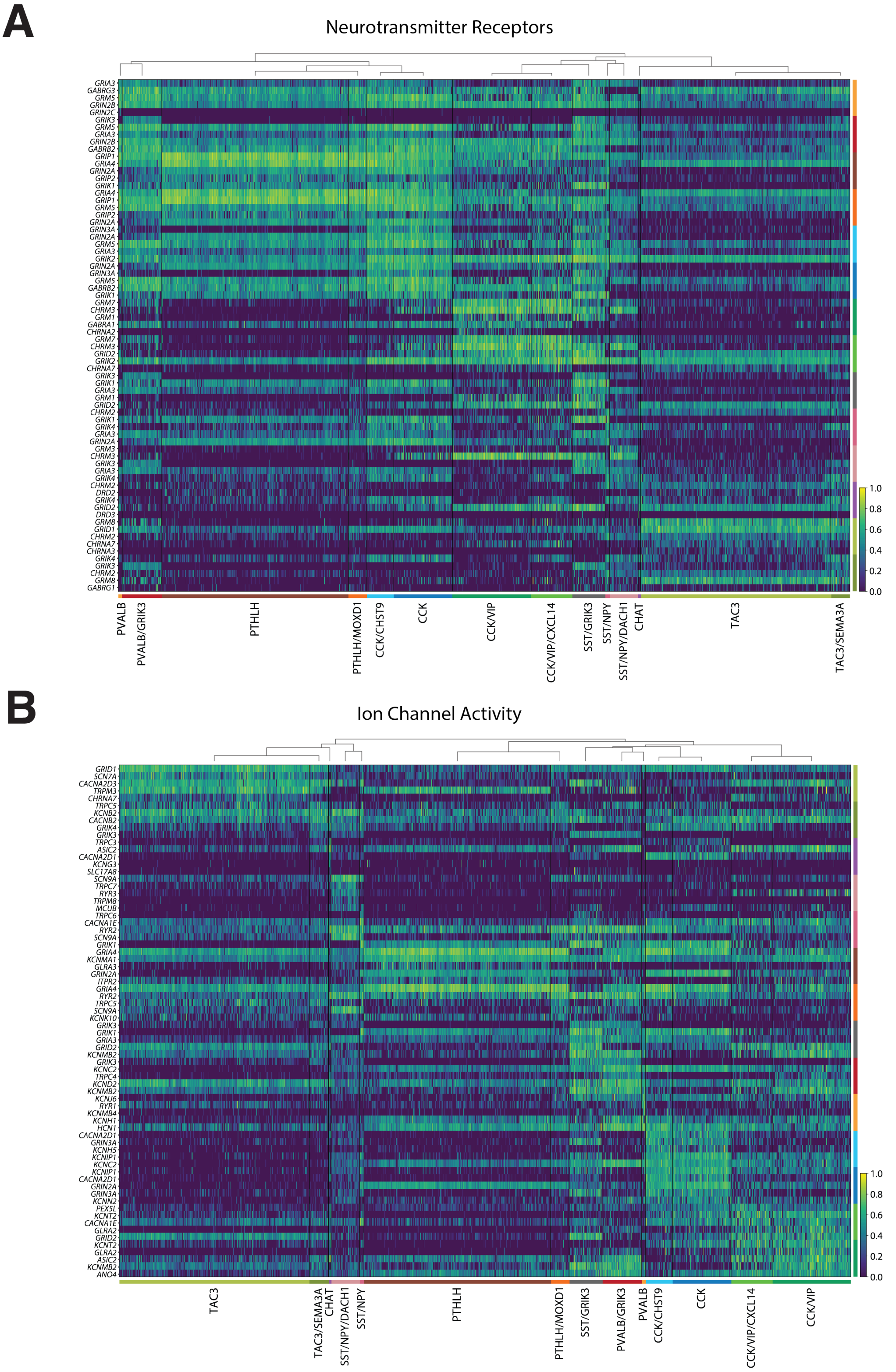


**Supplementary Fig. 5. A**. Heatmap depicting the top five differentially expressed genes by interneuron subclass among the neurotransmitter receptor genes. The dendrogram on top of the plot was computed based on the average Pearson correlation coefficient between groups across the selected subset of genes. **B.** heatmap including the top five differentially expressed genes by subclass in the ion channel subset. The dendrogram is computed as in A but considering only the subset of genes with ion channel activity in this case.

### Supplementary Figure 6

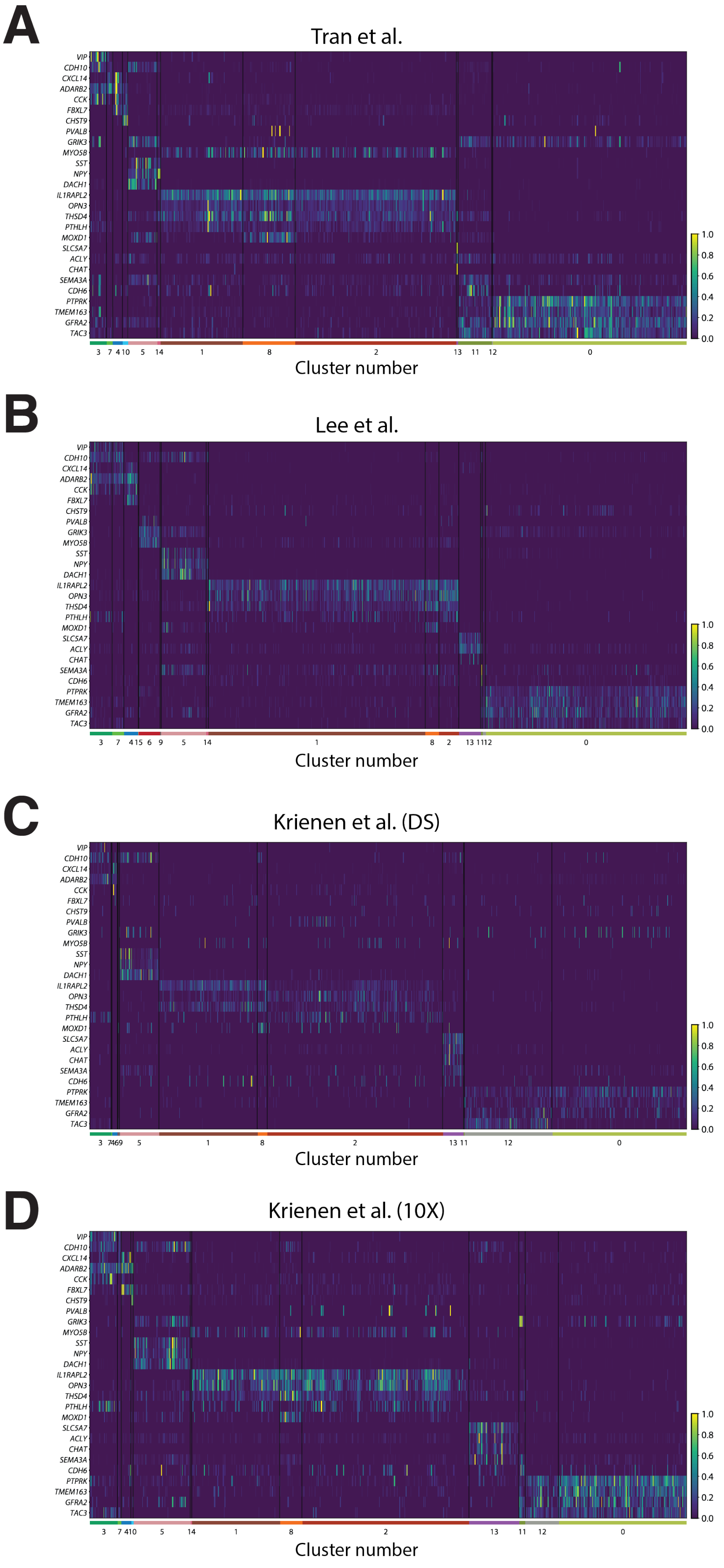

**Supplementary Fig. 6.** Expression of interneuron subclass markers on the raw counts of each of the public datasets used in the study. Each panel is labeled after the origin of the data. The raw counts of each gene are normalized by the maximum value of counts for that gene within each dataset (i.e. per row). The cluster numbers correspond to those on Figure 6, and the expression patterns follow the relationship established with the Interneuron subclasses.
